# Ungulate Responses to Unmanned Aerial Vehicles Flying at Different Altitudes in Africa’s Arid Savanna

**DOI:** 10.1101/2020.07.14.202093

**Authors:** Marlice vanVuuren, Rudie vanVuuren, Larry M. Silverberg, Joe Manning, Krishna Pacifici, Werner Dorgeloh, Jennifer Campbell

## Abstract

This paper tests the hypothesis that ungulate-UAV interaction depends strongly on flight altitude, that there may be a lowest altitude range for which the ungulates are not exceedingly disturbed, dictating a practically achievable level of discernibility in flight observation. This question strongly influences the future viability of the UAV in the study and protection of the ungulates in Africa’s arid savanna. This paper examined the behavioral responses of a group of free ranging ungulate species (Oryx, Kudu, Springbok, Giraffe, Eland, Hartebeest, and Impala) found in an animal reserve in Namibia to the presence of different in-flight UAV models. The study included 99 flights (337 passes) at altitudes ranging from 15 to 55 meters. The ungulates were unhabituated to the UAVs and the study was conducted in the presence of stress-inducing events that occur naturally in the environment. The results suggest strong correlations between flight altitude and response across the different ungulates and anecdotal evidence suggests in some cases rapid habituation to the UAVs.

## Introduction

The highly valued ungulates in Africa’s arid savanna, making up more than 25% of its landscape, might be effectively monitored by unmanned aerial vehicles (UAV) [1]. The question of this paper concerns the extent to which ungulate responses might allow aerial wildlife monitoring (AWM) by UAVs. The hypothesis is that ungulate-UAV interaction depends strongly on flight altitude, that flying too low could excessively disturb them, and that there may be a lowest altitude range for which the ungulates are not exceedingly disturbed (putting aside for the moment how exceedingly disturbed is quantified) – dictating some achievable level of discernibility in flight observation. This question is valuable because it strongly influences the future viability of the UAV in the study and protection of the ungulates in Africa’s arid savanna.

Several papers have already examined the effect of UAV altitude on non-ungulate response (data containing 5 to 100 trials). Biologists found that bears have an increased heart rate when the UAV flies 200 meters from the bear, 20 meters above it [2]. Scientists surveyed elephants in Burkina Faso, flying at altitudes of 100 meters and 300 meters over 10 km transects [3]. Researchers found that the influence of the UAV on the penguin is significant at an altitude of twenty meters [4].

Note that the AWM engineer or designer, when laying out a system of UAVs to monitor a field, would begin by selecting a flight altitude after which vehicles and a communication network would be selected, and in more advanced systems sophisticated data patching methods and graphical interfaces employed [1]. Other preparations, too, would be necessary depending on the application. Indeed, the large number of reported studies and methods preparing for AWM strongly suggests that the potential AWM applications are plentiful. Considering just a few of the papers, studies have focused on the cost savings [5], the management role [1], and more applied work on wildlife tracking [6], counting methods [7-9], and anti-poaching [10].

In general, the level of danger that an ungulate perceives strongly influences its response. The perceived danger that a UAV poses could come from the ungulate-UAV distance, sensed by an auditory signal, or it could come from secondarily observing the responses of neighboring species who have already responded to the UAV. In Africa’s arid savanna, one finds many ungulates in mixed herds so one could expect neighboring herds to trigger a response. The usual source of the danger to the ungulate comes from the ground, not the air. Therefore, the level of response could be moderate or low depending on flight altitude, while differing from specie to specie. One also expects the responses to depend strongly on the environment. The time of day (before or after eating), the proximity of nearby animals posing a danger, and the passing by of vehicles or other stress inducing events would be expected to influence the responses.

The method section describes the study site, the wildlife species monitored, the data collection process, the aerial vehicles, and the data collected. The results section gives the ungulate responses, the occurrences of positive responses (no or little discernable reaction) versus altitude, and discusses the data and secondary factors. Finally, the paper summarizes the results and draws conclusions.

## Method

The purpose of this study was to assess ungulate responses to UAVs flying at different altitudes in Africa’s arid savanna. Several considerations were vital when setting up the study. First, the protection of ungulate wildlife from human interaction favors human interactions that are sufficiently unobtrusive such that it requires no or a minimal level of habituation. Furthermore, ungulate habituation, when none of the conditioning is negative, would only tend to decrease the altitude that an ungulate tolerates. Therefore, for the purposes of this study, it was reasonable to focus on unhabituated ungulates, recognizing how the results of this study would extend to habituated ungulates. Secondly, in order for the results to apply to the natural habitat of the arid savanna, it was also important to perform the study in the mixed herd environment in the presence of its naturally occurring stress inducing events. The variable conditions of the natural environment introduce noise into the study that increases the difficulty to discern the effect of altitude on the response data. However, it can also determine whether or not UAV altitude is a dominant factor against the other factors, as hypothesized.

The ungulate response considered positive (null) or negative and distinguished between two levels of positive and negative responses to test out this hypothesis. The two levels of positive response were no movement (no discernable movement) and alert (eye/head movement and/or turning of the upper body). The two levels of negative response were relocation (moving out of the way of the UAV but still in sight) and flee (movement from sight).

The study was conducted at a wildlife sanctuary (N/a’an ku se) located about 40 km east of Windhoek, Namibia’s capital city, over a ten-week period in 2017 from September to November. The land is 25 km^2^, located on Namibia’s central plateau, and its elevation is 1600 to 1800 meters. From September to November, the average temperature is 20 to 23 C, and the average rainfall is 10 mm in September, 10 mm in October, and 30 mm in November. The land is a semi-closed eco-system surrounded by an 8-foot game fence that includes free ranging leopards and cheetah and the species are unhabituated to UAVs. Fig. 1 is a map of the test site. The study species included a group of ungulates commonly found in arid savannah eco-systems throughout Africa: Oryx (*Oryx gazella*), Eland (*Taurotragus oryx*), Giraffe (*camelopardalis*), Springbok (*Antidorcas marsupialis*), Hartebeest (*Alcelaphus buselaphus*), Plains Zebra (*Equus quagga*), and Kudu (*Tragelaphus strepsiceros*).

**Fig 1:**
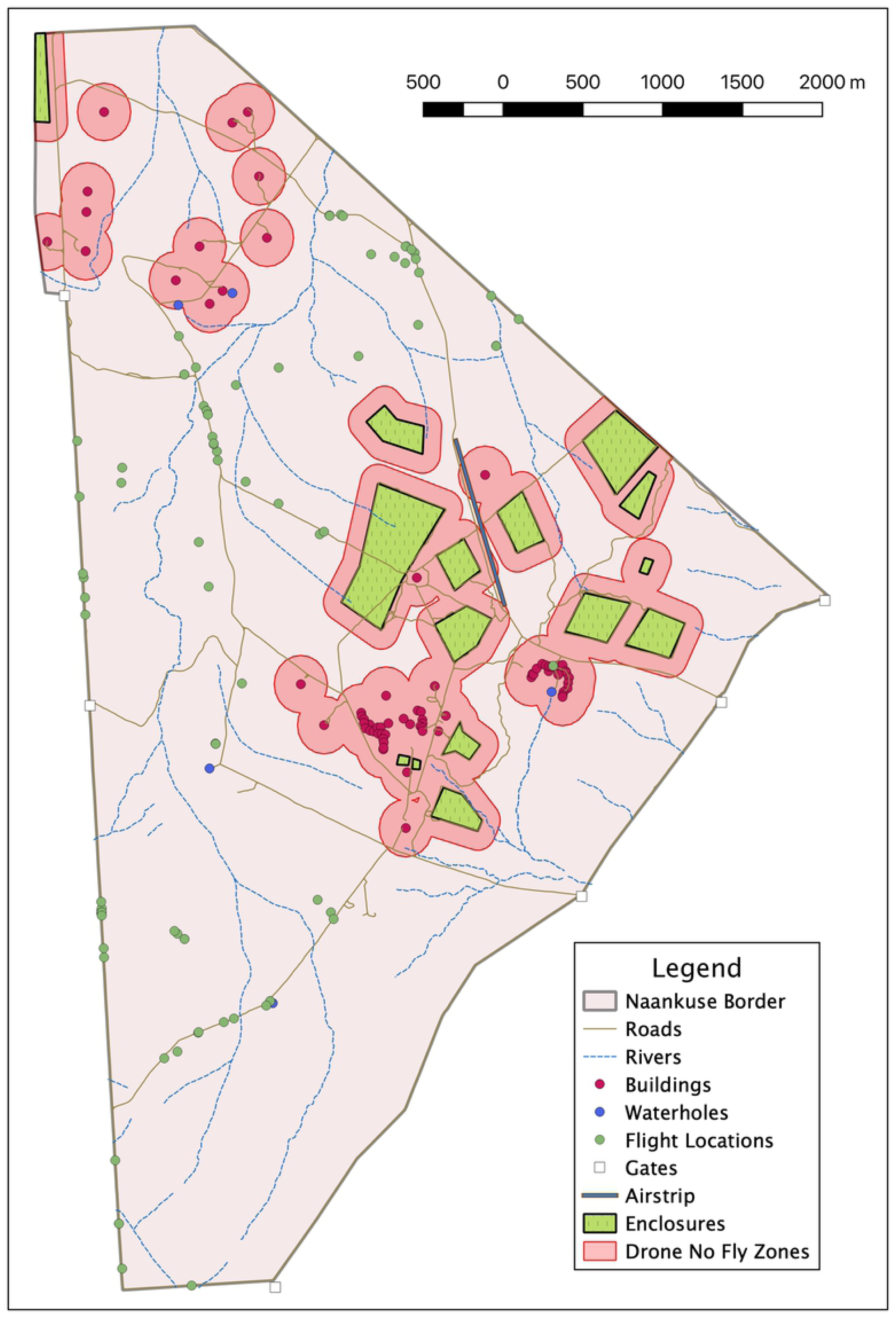
N/a’an ku sȇ Map and Flight Locations.

A field unit of 12 students collected data over a 10 week period (See the acknowledgement section). The field unit formed data collecting sub-units, each consisting of four personnel: (1) a pilot, (2) a primary observer, (3) a ground camera operator, and (4) a data recorder. The pilot was responsible for flying the UAV. When flying in RC mode, the pilot was also responsible for operating the flight camera and maintaining visual contact with the study species from the air. The primary observer was responsible for observing the species and maintaining visual contact with the ground station. The primary observer worked next to the data recorder and relayed all of his wildlife response observations throughout the flight to the recorder. The ground camera operator was responsible for photographing/video recording the responses. The data recorder was responsible for annotating information relayed from the primary animal observer on a flight-data collection sheet.

The suite of UAVs used consisted of the DJI Phantom 3 (Baseline), DJI Mavic Pro, Mavic Pro, Custom X8, and Skyeye. Table 1 shows the tested noise levels of the different UAVs with their baseline equivalent altitudes.

**Table 1:** UAV Noise Levels.

| UAV type | Frame size<br>(mm) | Weight<br>(g) | Noise level<br>(dBA) | Altitude<br>(m) |
| --- | --- | --- | --- | --- |
| Phantom 3 | 350 | 1280 | 60.24 | 50 |
|  |  |  | Baseline altitude | 50 |
| Mavic Pro | 335 | 734 | 45.7 | 50 |
|  |  |  | Phantom Baseline equivalent altitude | 266.7 |
| Skyeye | 2000 | 2000 | 52.5 | 50 |
|  |  |  | Phantom Baseline equivalent altitude | 121.9 |
| Custom X8 | 650 | 4400 | 65.0 | 50 |
|  |  |  | Phantom Baseline equivalent altitude | 28.9 |

As shown, the loudest to quietest vehicles were the Custom X8, Phantom 3, Skyeye and Mavic Pro. Noise disturbance is not only influenced by the noise which can be qualified by sound intensity (dB) and frequency (Hz), but also by its duration and pattern. The age and physiological state of the animal at the time of exposure, the exposure history of the animal, and the predictability of the acoustic stimulus can also play a role [11]. The effect of altitude on animal response was studied in the presence of all of these naturally occurring factors.

The design of the experiment began with determining a range of flight altitudes. The range of flight altitudes was determined through 24 preliminary flights, flying at altitudes from 20 meters to 100 meters in 10-meter increments. We determined that 15 meters was a suitable lower bound because of safety concerns and that 55 meters was a suitable upper bound because negative response had already significantly dropped off by that altitude. The data collection process began every morning with locating animal herds and setting up a field unit nearby. Local personnel understood the best locations to set up field units and launch the UAVs. As shown in Fig. 1, the launch points were located in the low-lying, open areas along the roads. The data collecting personnel hiked or travelled by ground vehicle to those launch points.

The horizontal distances between the animals and the field unit points varied widely – between 50 and 600 meters (measured in 25 m increments). The units logged this data to determine its effect on the responses, if any. The total data set consisted of 99 flights (337 passes) that ranged in altitude from 15 to 55 meters. Each pass began with taking off from a launch point, flying about 200 m away from a spotted animal group, changing altitude as appropriate, and then flying 400 m, passing over the animal group. The UAV was flown over the animal or herd at 10 m/s. The next pass was flown at another randomly selected altitude. If an affirmative response was invoked (relocation or flee), the drone would wait at a distance while the animals would settle back to a sedentary behavior. Passes were continued until animals were no longer within sight. Table 2 shows the procedure that each of the members of the field unit followed.

**Table 2:** Field Unit Procedures.

|  |  |
| --- | --- |
| Pilot | Pre-flight checklist, flight, post-flight checklist, reports damage or concerns at end of day |
| Camera operator | Pre-flight and post-flight checklist for camera and gimbal, ensure data is downloaded, sd card cleared at end of day, assists pilot with DJI drones. |
| Animal observer | Observes surrounding field prior to takeoff for safety, observes animal responses during flight, decides whether to continue or end mission based on animal behavior. |
| General observer | Ensures all of the roles are being properly executed, ensures that no wildlife is encroaching upon during the flights, assists the data collector in obtaining data during the flights when several tasks are being performed simultaneously. |
| Data entry | Enters all of the pre, during, and post flight data |

## Results and Discussion

### Species Reactions

The relative abundance of the different species varies from site to site and by season. Table 3 shows the populations of the monitored species at the test site. In some cases, single specie herds were observed while in other cases there were mixed herds.

**Table 3:** Number of passes (% of total passes)

| Animal | Single Species | Mixed Species |
| --- | --- | --- |
| Oryx | 63 (15.79%) | 110 (27.57%) |
| Zebra | 24 (6.02%) | 95 (23.81%) |
| Kudu | 2 (0.5%) | 4 (1%) |
| Springbok | 40 (10.03%) | 101 (25.31%) |
| Giraffe | 49 (12.28%) | 58 (14.54%) |
| Eland | 23 (5.76%) | 54 (13.53%) |
| Hartebeest | 20 (5.01%) | 32 (8.02%) |
| Impala | 0 (0%) | 3 (0.75%) |

**Table 3:** Response data for giraffe over time.

| Sets of 15 passes | First | Second | Third | Fourth |
| --- | --- | --- | --- | --- |
| No. of Negative Responses | 4 | 3 | 3 | 2 |

As shown, the least populated species were the Impala and the Kuda and the most populated were the Oryx, Zebra, Springbok and the Giraffe. Also notice that the Oryx, Zebra and Springbok were most frequently found in a mixed herd and that the Giraffe were found in a mixed herd about half the time.

Figs. 2 and 3 show the different ungulate responses for all the single species flights and the mixed species flights. We did not show the Kudu and Eland responses because the frequency of their sightings was too low. There was strong similarity in the responses across animal type and across single species versus mixed species. The positive responses for each ungulate occurred between 68% and 95% of the time. The lowest negative (affirmative) response was the Giraffe at 10% and the highest negative response was the Zebra at 32%.

**Fig 2:**
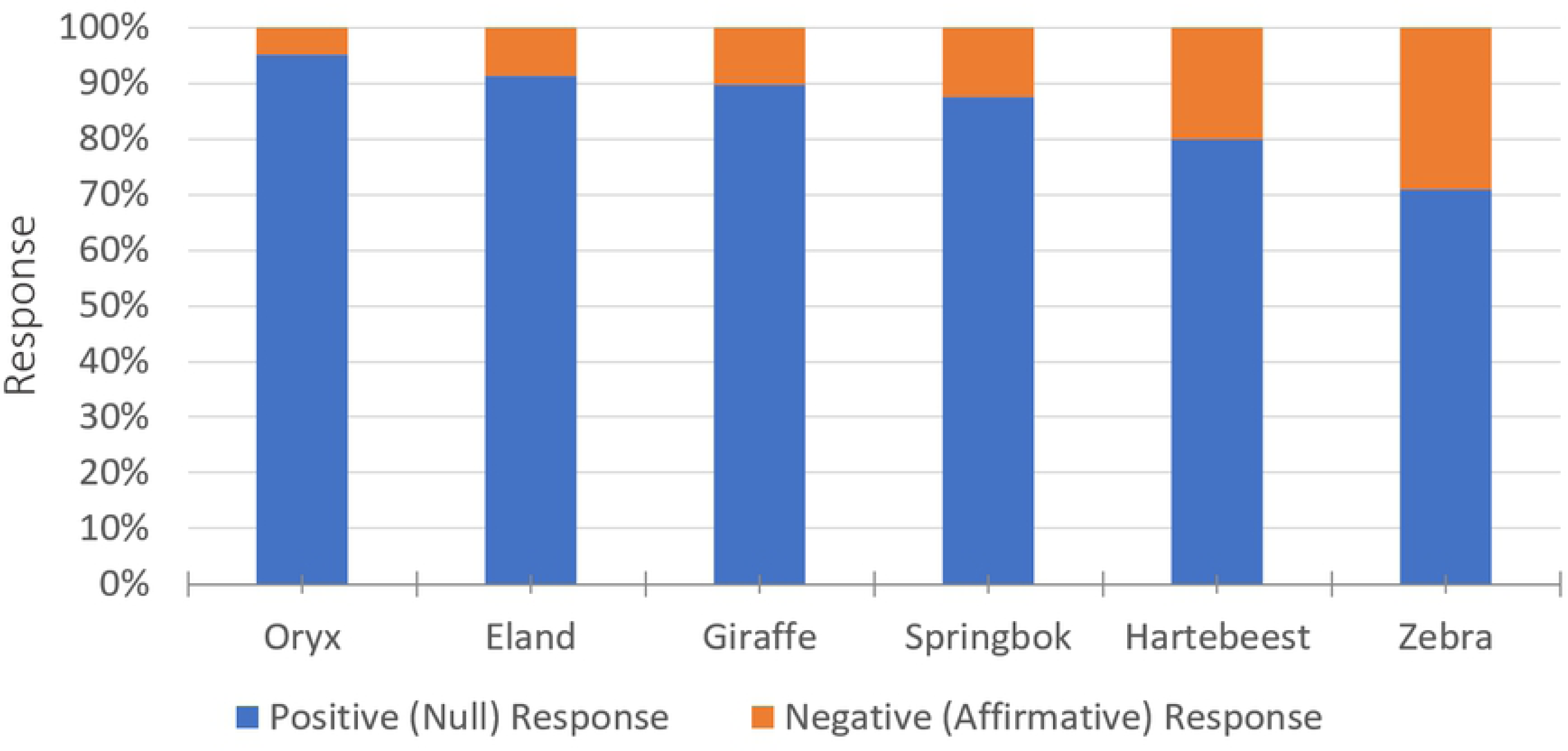
Normalized Single Species Responses.

**Fig 3:**
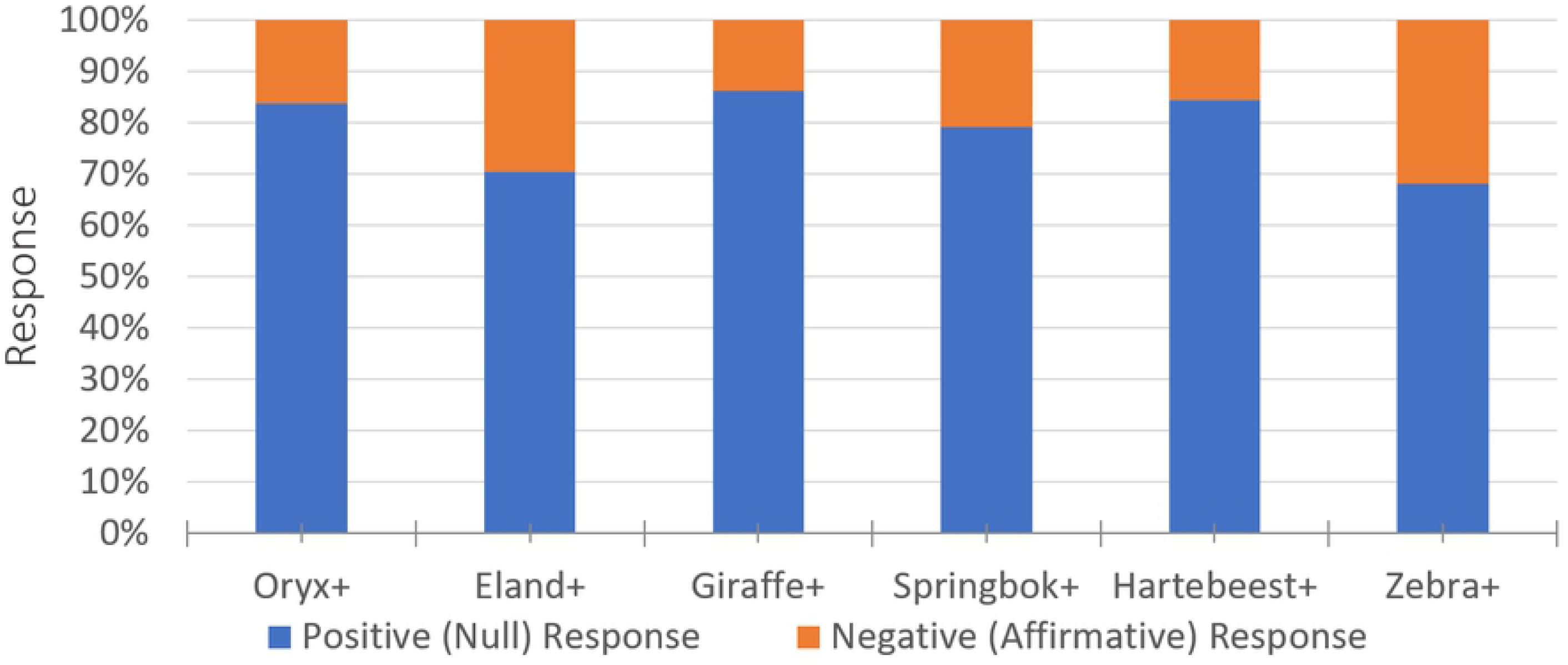
Normalized Mixed Species Responses.

The species responses to UAV shown above depended on a variety of factors related to the danger the UAV posed in relation to other present dangers. The field unit observed in individual cases the effect of their proximity to the UAV, the time of day, and whether or not the UAV appeared to be competing with attention of a larger source of danger. Vehicles that drive by, nearby baboon/cheetah walks, and weather all appeared to influence the responses, as well. General species behavioral patterns influenced the results, too. Generally, most ungulates graze or browse in early morning and in early evening/late afternoon, allowing them to avoid the heat during midday, during which they hide from predators under the shade of trees. Some ungulates are browsers (eat leaves) and are found in thick bush, so they are more difficult to spot and observe. Others are grazers, found in open grassland, and therefore are easier to spot. Impala and Kudu numbers were lower simply because fewer of them are free ranging on the property. Kudu are very cryptic/elusive animals, often hiding in bush cover in small herds. Most of the data were collected from larger open areas where the animals were easier to locate and where they could be observed from a distance without disturbing them prior to take-off. We noted differential responses for some species when in single species groups versus in mixed species groups. For example, the Zebra are flighty when not with other ungulates but appeared to react just like the Oryx when with them. The Giraffe were observed to be flighty when first passing over them but became habituated to the UAV presence after just two passes.

More generally, pertaining to animal habituation to humans and traffic, we used vehicles to transport ourselves and the vehicle traffic appeared to affect the responses. We observed the animals to be more flighty in areas that are closer to more traveled roads and it was difficult to collect data close to neighboring farms that allow game hunting since the animals there were flighty when sensing human presence. Male responses may have also differed from female responses. For example, some male springboks may have suppressed their flight responses to keep hold of their territory while the females and maternity herds exhibited a flight response.

### Visual and Auditory Influence

Fig. 4 displays the percent makeup of each response for the given altitudes. First notice that the No movement response (in blue) increases with altitude, that the relocation and flee responses (grey and yellow) decrease with altitude, and that in between the alert response (orange) does not show a significant increasing or decreasing trend. Next notice at 15 meters that there was a positive response (No movement and Alert) 60% of the passes and that at 55 meters there was a positive response 90% of the passes. This figure helps illustrate the general pattern of less negative response at higher altitudes supporting the hypothesis that altitude plays a large role in animal response from UAVs.

**Fig 4:**
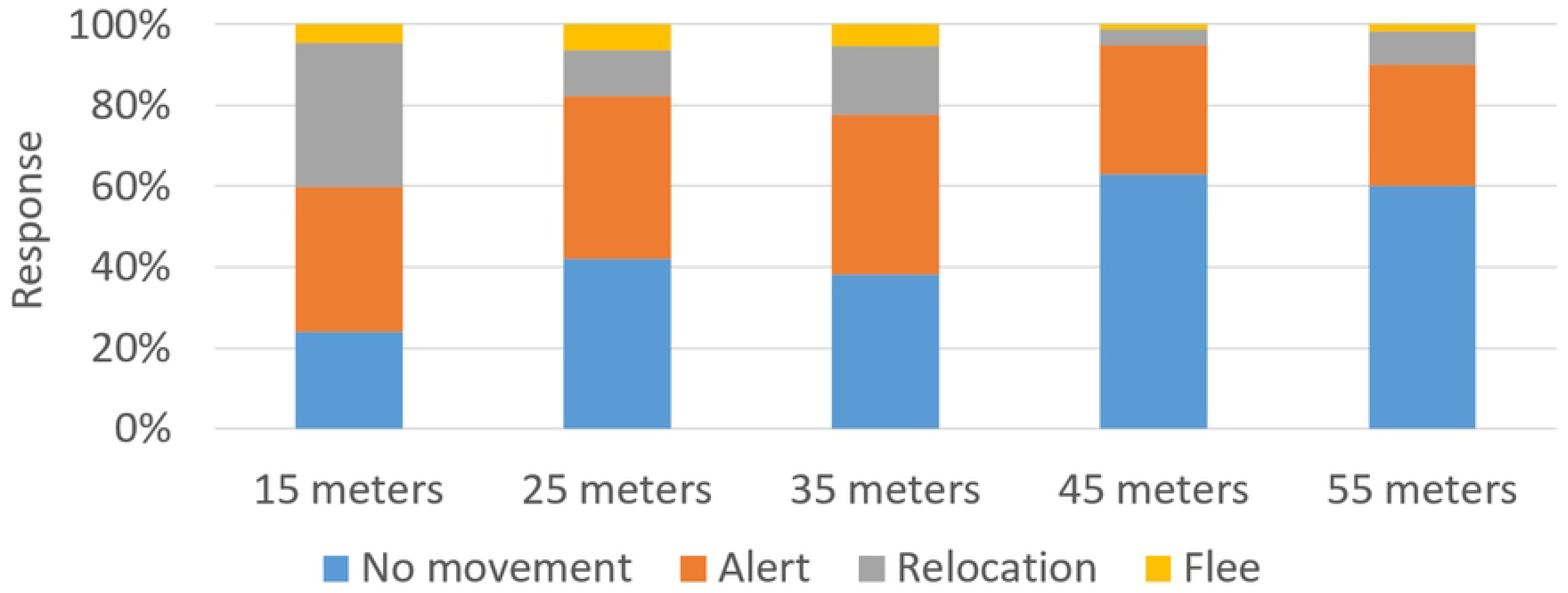
Combined Vehicle Response.

Two other visual factors were examined to more closely consider the impact of the visual role the UAV may play. These two factors were type of UAV, or flight mechanism, and apparent size in the sky. To examine the apparent size in the sky, the largest dimension was used and the altitude during the pass to create an estimated field of view (FOV) that the UAV would take up when looking up at it. The largest vehicle was the Skyeye fixed wing and the smallest was the Mavic Pro. No apparent trend in the data was found. Any trend was largely suppressed because the Skyeye had minimal disturbance, only invoking a negative response 1 time out of the 21 flights it made. It was the largest vehicle by far giving very low percent response at large FOVs. This mixed with the fact most of the other low percent affirmative data points were small FOVs gave the data no distinguishable trend. This however leads to the fact that the other visual factor, flight mechanism, does seem to play a considerable role. The fixed-wing data showed a considerably lower percentage of response than the multirotor data. This suggests that flight mechanism may impact animal response. A fixed wing looks more like a flying bird that does not pose a threat to an ungulate than an unfamiliar object with large spinning propellers. Fig. 5 shows the combined data averaged across the different vehicles with the error bars displaying the 95 percent confidence interval.

**Fig 5:**
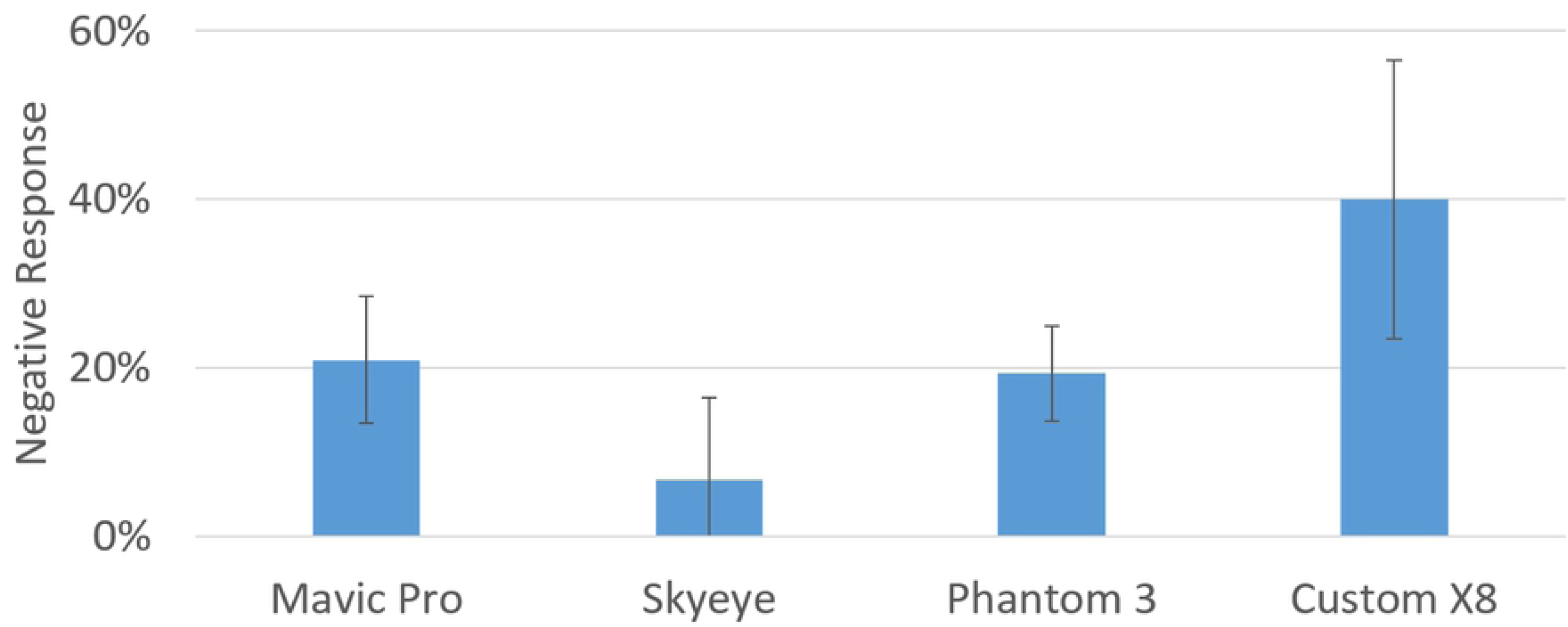
Response vs Vehicle.

Fig. 5 also gives insight into the general trend of animal response to sound intensity. The relative sound intensity each UAV produces was previously given in Table 1, the least noisy being the Mavic Pro and the loudest being the Custom X8 shown in order on Fig. 5 from left to right. If sound intensity were the driving factor, the Mavic Pro would have the lowest negative response percentage. However, it has nearly the same negative response percentage as the Phantom 3. Fig. 6 shows the Mavic and Phantom negative response plotted against sound intensity data and altitude for comparison.

**Fig 6:**
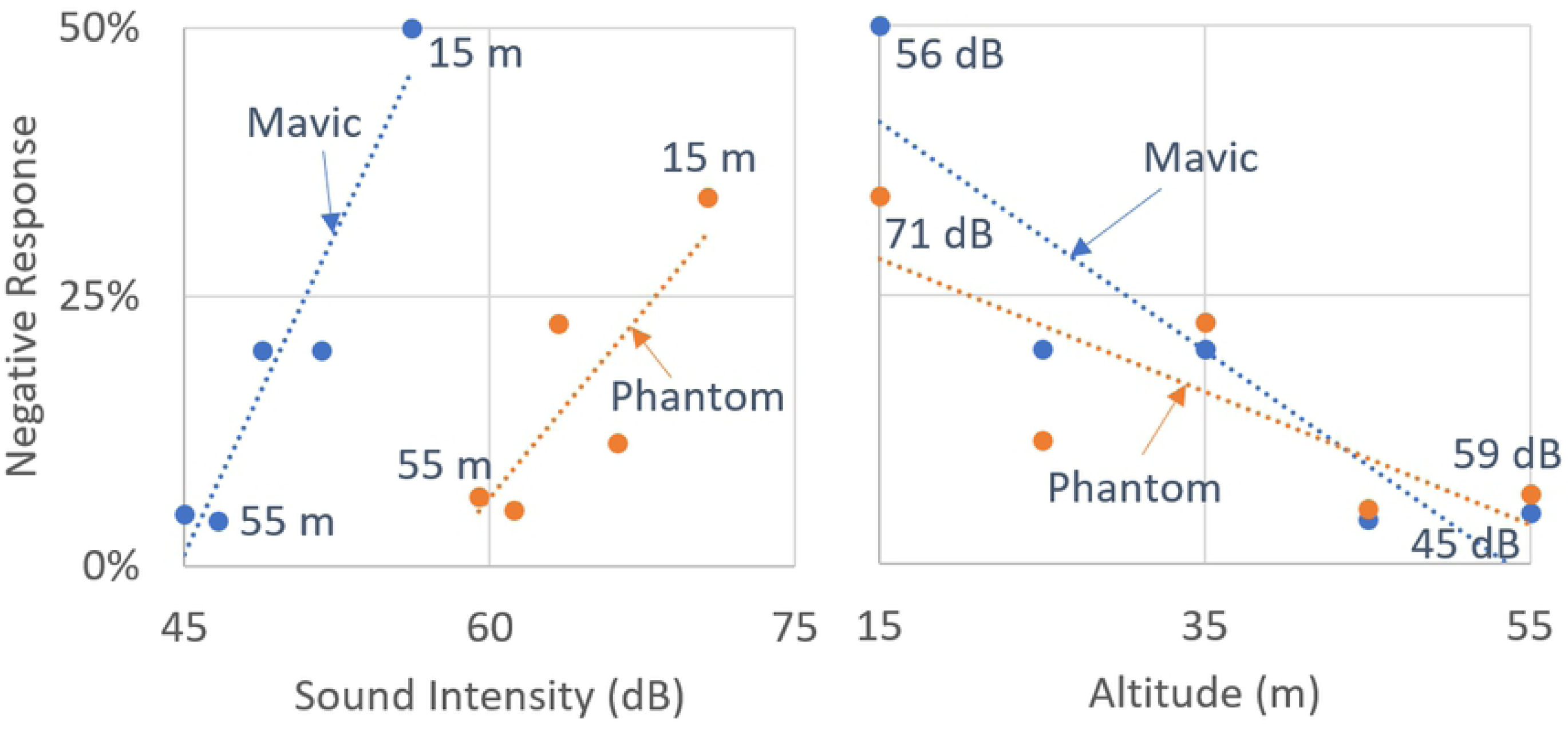
Negative Response vs Sound Intensity and Altitude for Phantom and Mavic.

In Fig. 6, the graph on the left, negative response vs sound intensity, shows a good linear representation of the two data sets (R^2^ = 0.91 for Mavic and R^2^ = 0.69 for Phantom). However, they have a large separation between them, a roughly 15 dB offset of the datasets. Now looking at the graph on the right, negative response vs altitude, shows the same datasets plotted with altitude. The most important takeaway being that the datasets overlap and almost align when plotted against altitude. This shows a stronger correlation to altitude than sound intensity for these two vehicles. This was also reinforced after taking the Pearson correlation coefficient for the two data sets. The coefficient was 0.303 for the negative response vs sound intensity for the combined vehicle data.

The coefficient was -0.836 for the negative response vs altitude. This shows a stronger correlation to negative response vs altitude for these two vehicles. In Fig. 7, the graphs show the Phantom 3 and Custom X8 plots with negative response vs sound intensity on the left and negative response vs altitude on the right.

**Fig 7:**
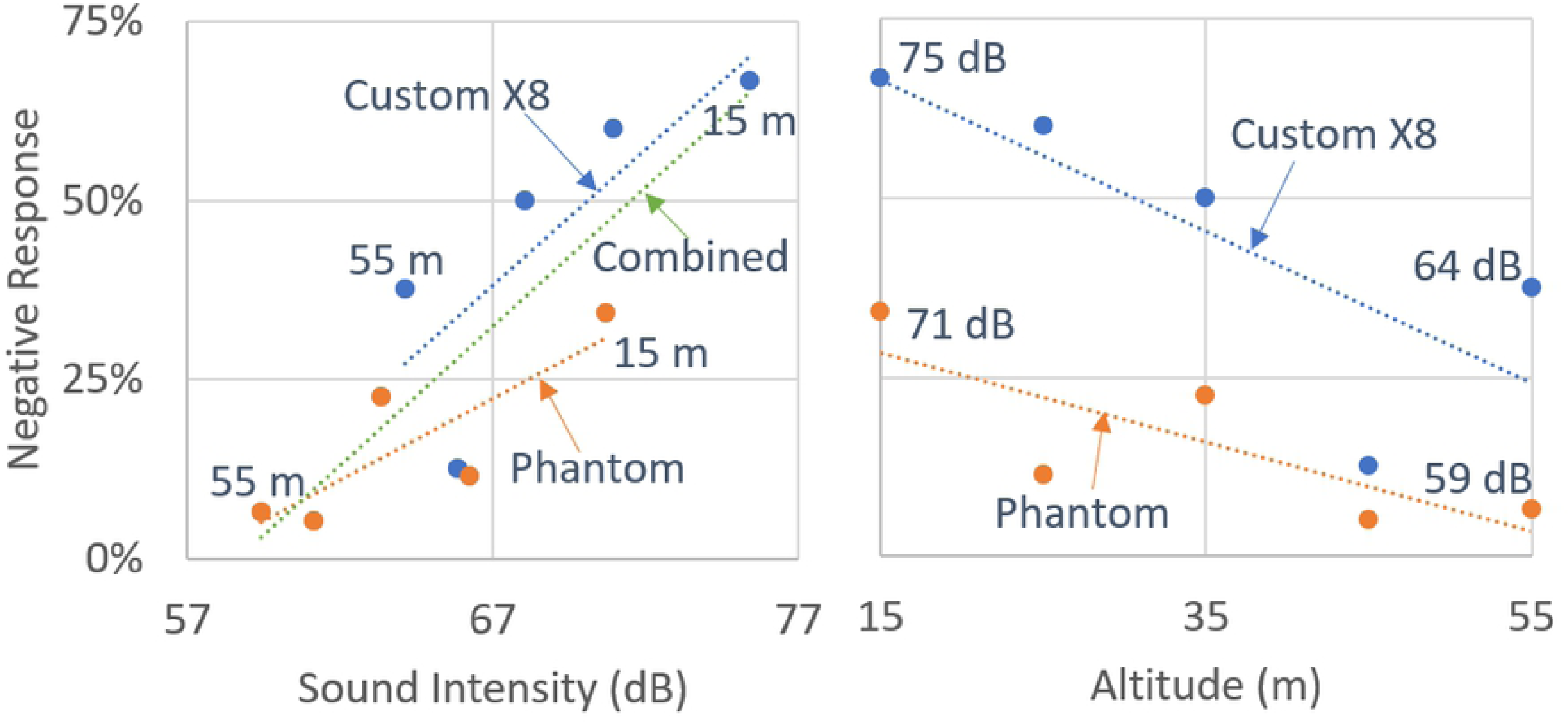
Negative Response vs Sound Intensity and Altitude for Phantom and Custom X8.

Unlike the fig. 6 where the data points overlap for the Mavic Pro and Phantom 3 when plotting the negative response vs altitude, the data points for the Custom X8 and Phantom 3 show an offset shifted in the direction of negative response. However, when the two UAVs are plotted showing negative response vs sound intensity, Fig.7 on the left, they nearly overlap. A trendline was plotted in fig. 7 of the combined dataset to show the linear relationship (R^2^ = 0.70). Similarly, the Pearson correlation coefficient of the was calculated for both negative response vs sound intensity and altitude. The coefficient was 0.837 for the negative response vs sound intensity for the combined vehicle data. The coefficient was -0.553 for the negative response vs altitude. This shows a stronger correlation to negative response vs sound intensity for these two vehicles. These relationships are inconsistent with the comparison of the Mavic Pro and Phantom 3.

An important thing to notice is that the slopes of the trendlines for the negative response vs altitude of the Phantom 3 and Custom X8 area roughly the same. The negative response of the Custom X8 is just shifted at each altitude of approximately 25% higher rate of negative response. One theory is there is a sound intensity threshold for which the animals begin responding more to the noise than the proximity of the UAV. This would explain why the Custom X8, being much louder than the other vehicles, exhibits a much higher negative response at each altitude. But when the vehicles are under the sound intensity threshold, like the Mavic Pro and Phantom 3, the vehicles more closely follow a response with respect to altitude. One final note on visual and auditory influence, is that when the higher the UAV flies and quieter it is near the animals the less likely it is to induce a negative response. Of the 55 passes flown at both above 50 meters and at or below 60 dB, only 3 negative responses were induced, and they were all relocation.

### Habituation

UAV had never regularly flown over the property where this research was conducted. Therefore, the research applies unhabituated ungulates. Of course, one would expect the percentage of positive responses to increase if the species were to become habituated to the presence of UAVs. The property contained 13 giraffe who stayed in a single herd. During the preliminary flights, we passed over the giraffe twice and each time exhibited a flee response. In the subsequent research we passed over them 60 more times and they never exhibited a flee response. This suggested that the giraffe might have been rapidly habituated to the presence of the UAVs warranting further study of the giraffe responses. We divided the data chronologically into 4 sets each containing 15 passes. The altitudes of the passes over each set were about 35 meters. Table 3 shows the negative responses over each set of passes.

The period in which this data was collected was 41 days. The 4 sets of passes varied slightly in time, having collected the first set in 11 days, the second set in 17 days, the third set in 8 days, and the fourth set in 5 days. While these numbers are statistically insignificant, there appears to be a trend toward less response over more passes. Although beyond the scope of this study, this anecdotal data suggests that animal habituation to the presence of UAVs is possible.

## Summary and Conclusions

This paper reports on the behavioral responses of free ranging ungulate species (Oryx, Kudu, Springbok, Giraffe, Eland, Hartebeest, and Impala) housed in an animal reserve in Namibia to the presence of different UAV models. The study included 99 flights (337 passes) at altitudes ranging from 15 m to 55 m. The results suggest strong correlations between flight altitude and animal response across the different ungulates and anecdotal evidence suggests rapid habituation to the UAVs. Over the suite of UAVs used in the study, the correlation between negative response and flight altitude was most pronounced at low sound intensities (less than 60 dB). At higher sound intensities (above 60 dB), it was found that the negative response has a more discernable parallel to sound intensity.

The single herd and mixed herd observations conducted in the study provide evidence in support of the feasibility of aerial wildlife monitoring (AWM) by UAVs. The highest percentage of positive responses occurred at the highest altitudes with UAVs having the lowest sound intensity, finding only a 5% chance of inducing a negative response when flying at 55 meters and when using a UAV that is quieter than 60 dB. The study was conducted with ungulates that were unhabituated to the presence of UAVs. Further study of habituation to the presence of UAVs could further strengthen the results.

## Acknowledgments

The faculty advisers and Office of Global Engagement staff at North Carolina State University made this study possible. Namibian personnel and authorities were instrumental, too, as were the previously unnamed students: Hussain Arif, Kennedy Brinson, John Dvorak, Joel Forsyth, Josh Glazer, Jacob Keller, Graham Lutz, James Ploss, Allison Schumacher, Nicholas Skahill, and Leena Vo.

